# A small increase in *CHEK1* activity leads to the arrest of the first zygotic division in human

**DOI:** 10.1101/2020.12.26.424381

**Authors:** Beili Chen, Jianying Guo, Ting Wang, Qianhui Lee, Jia Ming, Fangfang Ding, Haitao Li, Zhiguo Zhang, Lin Li, Yunxia Cao, Jie Na

## Abstract

The first mitotic division in mammalian zygotes is unique. The fertilized egg reactivates its cell cycle, and the maternal and paternal genomes start to reprogram to become totipotent. The first division is very sensitive to a range of perturbations, particularly the DNA damage, leading to the embryo's failure to enter the first mitosis. We discovered that a point mutation in the human *CHEK1* gene resulted in an Arginine 442 to Glutamine change at the C-terminus of the CHEK1 protein. *CHEK1* R442Q mutation caused the zygote to arrest just before the first division. Heterozygote individuals appeared to be healthy except that the female carriers are infertile. Expressing the corresponding mouse mutant Chk1 protein in zygotes also caused arrest before the first mitosis. Treating Chk1 R442Q mouse zygotes with low concentrations of CHEK1 inhibitor enabled the embryos to overcome the cell cycle arrest and resume normal development. Our results revealed an unexpected zygote mitotic checkpoint, which is extremely sensitive to the CHEK1 kinase activity. The fine-tuning of the DNA damage checkpoint permits the arrested one-cell embryos to overcome the first mitotic block and develop into healthy animals. These findings have important implications in assisted human reproduction.

---

During the first cell cycle, the mammalian zygotes switch from meiosis to mitosis, both paternal and maternal genomes replicate, and the minor genome activation occurs. In the meantime, the male and female pronucleus move to the center of the zygote and merge. Then zygotes enter the metaphase, and sister chromatids separate into two daughter cells (Clift and Schuh, 2013). Significant epigenetic reprogramming also takes place during this sensitive time window. Thus, many perturbances may cause the first mitosis to fail.

We discovered a family that the female adults were infertile. One female patient underwent in vitro fertilization treatment. Interestingly, 8 out of 15 eggs were fertilized normally as the zygotes contained one female and one male pronucleus, but they all failed to divide into 2-cell embryos (Table 1). Further investigation found that three sisters of this female patient were infertile, while another two sisters had children. With the patients’ consent, whole-exome sequencing was carried out in all of the infertile subjects. After the pedigree analysis, we found that a heterozygous missense variant in the *CHEK1* gene, c. G 1325A; p.R442Q, was shared by all the infertile sisters (Figure 1A). The variant was confirmed by Sanger sequencing in the whole family members (Figure 1B). The results suggested that each infertile subject inherited the variant from their father II-8, and the variant was segregated within the whole family (Figure 1A and 1B). CHEK1 protein plays an essential role in the DNA damage response (Chen et al., 2012; Ju et al., 2020; Liu et al., 2000) and is conserved across species (Supplementary Figure 1A). Interestingly, although the same *CHEK1* R442Q variant is expressed in somatic cells, based on RT-PCR and Sanger sequencing analysis of the peripheral blood samples from III-2,3,4,5,6 and 7 (Figure 1C and Supplementary Figure 1B-D), the heterozygote carriers are all healthy. We also analyzed the transcriptome profile of human oocytes, 1 and 2-cell embryos, *CHEK1* transcripts are present at high levels in GV and metaphase II oocytes, zygotes, and 2-cell embryos (Figure 1D). Therefore, it should exist as a maternal protein. Thus, we highly suspect that the *CHEK1* R442Q mutation caused the first zygotic mitosis to arrest.

**Table1.**
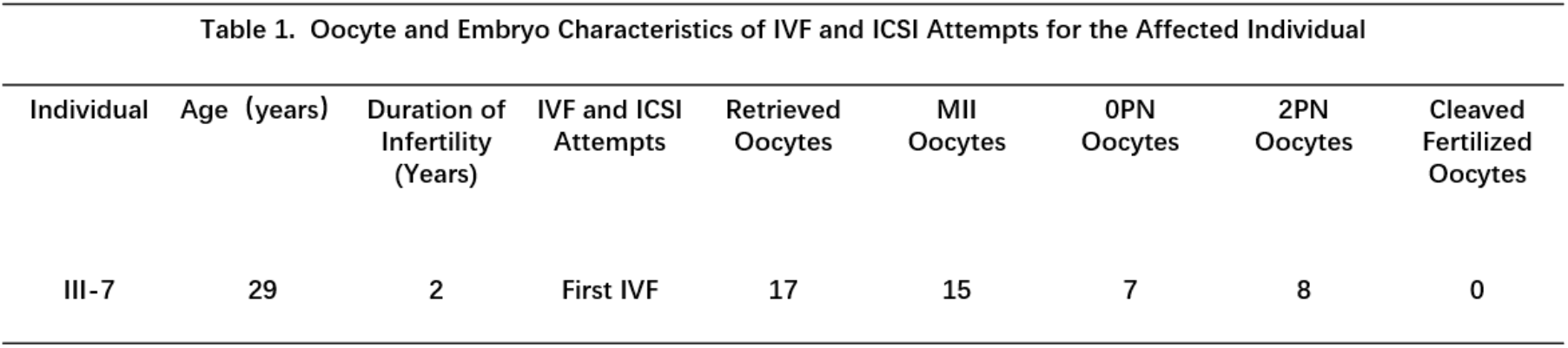
Oocyte and embryo phenotypes of IVF attempt for the *CHEK1* R442Q female patients.

**Figure 1.**
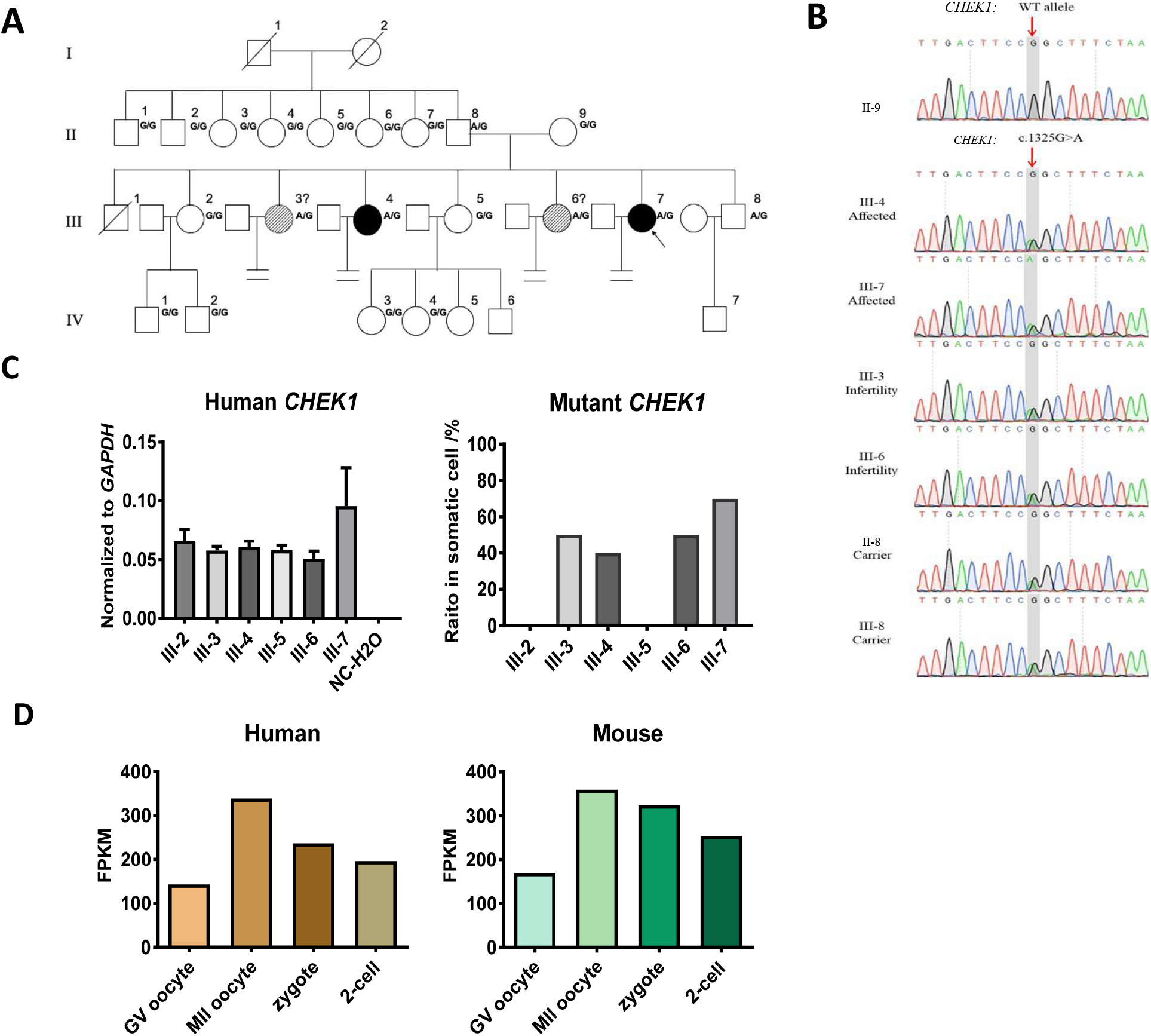
First mitosis failure in zygotes from *CHEK1* R442Q females. (A)Genetic analysis in the families carrying *CHEK1* R442Q. Black circles indicate the affected individuals, and the circles with slashes indicate the suspected individuals (III-4 and III-7 had been confirmed during IVF cycles, III-3 and III-6 females were infertile, possibly caused by the arrest of the embryo cleavage). The black arrowhead indicates the proband. (B)Sanger sequencing result of *CHEK1* R442Q mutation in Affected, Infertile, and Carrier individuals. Double peak at the 1325 position highlights the mutated nucleotide. (C)QPCR result confirmed that *CHEK1* R442Q transcripts are present in patients’ somatic cells. Left panel, the expression levels of CHEK1 mRNA in human blood cells. Right panel, the ratio of *CHEK1* R442Q mutant transcript in patient somatic cells verified by Sanger sequencing. (D)*CHEK1* expression levels in human and mouse GV oocyte to 2-cell embryo based on RNA-Sequencing data from GSE101571, GSE36552, and GSE71434.

To test this hypothesis, we cloned mouse Chk1, whose amino acid sequence is 93.07% identical to that of the human CHEK1, and generated the same Chk1 R442Q mutation as found in the infertile patient. Upon overexpression of WT or R442Q Chk1 protein in mouse zygotes by mRNA injection, we found that more than 60% of the Chk1 R442Q zygotes failed to divide into 2-cell embryos (Figure 2A and B). Overexpression of Chk1 WT did not affect the first mitosis (Figure 2B and C). It has been shown that overexpression of Chk1 in mouse GV oocytes caused meiosis I arrest (Chen et al., 2012). Interestingly, when we overexpressed Chk1 WT and R442Q in GV oocytes, the R442Q caused significantly more oocytes to arrest in metaphase I than WT (Figure 2D-F). We analyzed Chk1 protein localization based on the immunofluorescence signal, both Chk1 WT and R442Q proteins localized in the nucleus and on the spindle during mitosis and meiosis (Supplementary Figure 2A and C). The amount of overexpressed mutant protein is less than the overexpressed WT protein (Supplementary Figure 2B and D), based on their respective mean fluorescence intensity (MFI), suggesting that the R442Q change did not stabilize Chk1 protein.

**Figure 2.**
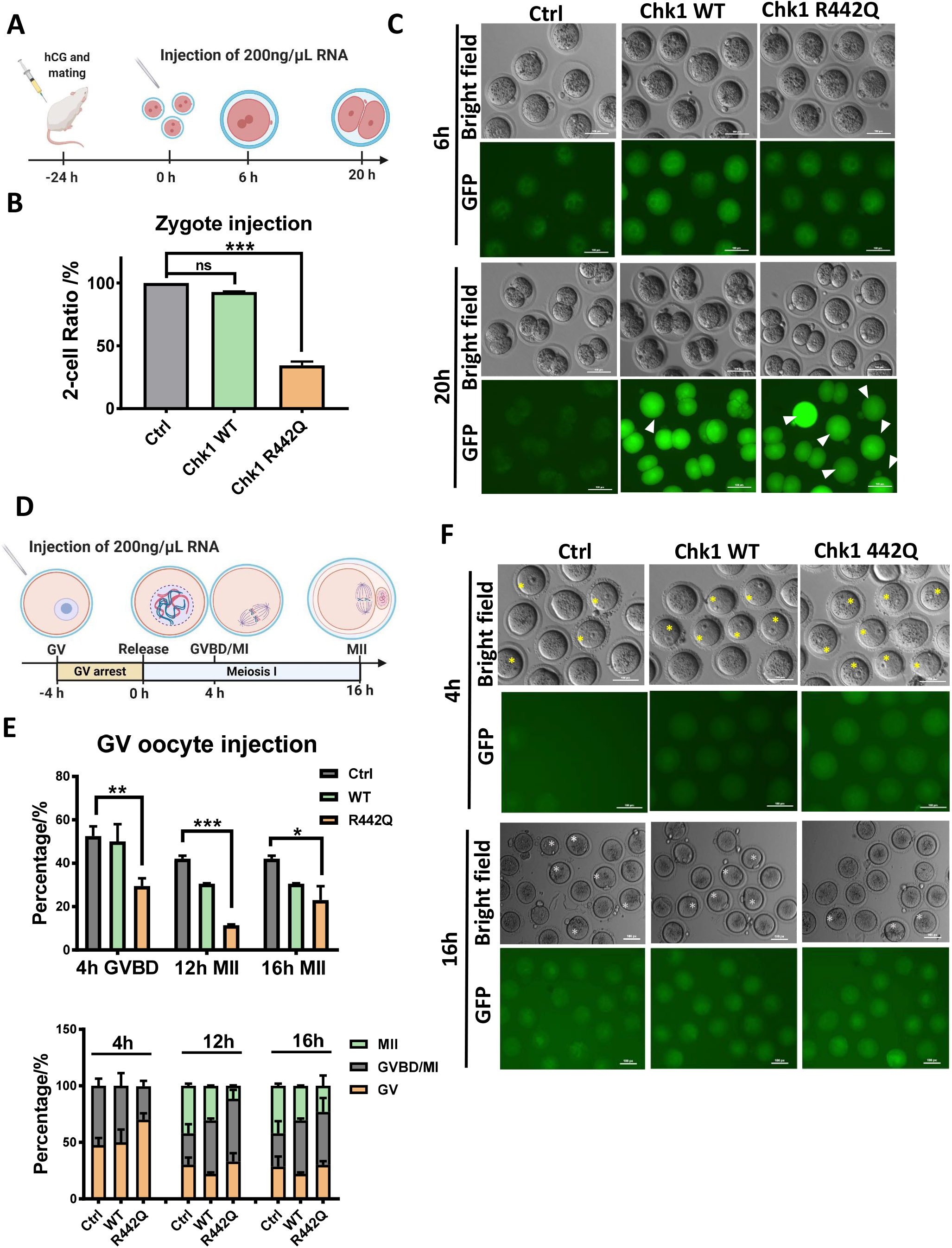
Overexpression of *Chk1* R442Q inhibited the first mitosis in mouse zygotes and meiosis in GV oocytes. (A)Schematic of overexpressing *Chk1* WT and R442Q in mouse zygotes by mRNA microinjection (Created with BioRender.com). (B)Bar plot showing Chk1 R442Q arrest the first cleavage. Ctrl, n=23. WT, n=48. R442Q, n=51. All pooled from 2 separate experiments; one-way analysis of variance (ANOVA), Bonferroni’s test for individual comparisons. Bars are means ± s.e.m. (C)Representative live cell images of zygotes microinjected with mRNA encoding Chk1 WT and R442Q GFP fusion protein. White arrowheads point to the arrested embryos. Time is as indicated. Scale bar, 100 μm. (D)Schematics of overexpressing Chk1 WT and R442Q in mouse oocytes (Created with BioRender.com). Microinjected GV oocytes were first blocked in the GV stage with Cilostamide for 4h. GVBD rate and MII rate 4h, 12h, 16h after release from blocking were counted. (E)Bar plot showing Chk1 R442Q delay mouse oocytes development. Ctrl, n=50. WT, n=59. R442Q, n=52. All pooled from 2 separate experiments. Bars are means ± s.e.m. Upper panel, statistical analysis of the percentage of GVBD 4h after release, MII 12h, and 16h after release. Two-way analysis of variance (ANOVA), Bonferroni’s test for individual comparisons. Lower panel, composite bar graph of percentages of GV, GVBD, and MII oocytes of Ctrl, Chk1 WT, and R442Q groups. (F)Representative live cell images of oocytes microinjected with mRNA encoding Chk1 WT and R442Q GFP fusion proteins. Yellow and white stars point out GV arrested oocytes (4h), and oocytes progressed to the MII stage (16h). Scale bar, 100 μm.

As the activation of CHEK1 kinase in response to DNA damage will cause cell cycle arrest in the G2 phase (Patil et al., 2013), we hypothesize that the R442Q mutation may cause an increase in CHEK1 protein kinase activity. Therefore, we generated the N-terminus kinase domain, the full-length, and the full-length R442Q mutant CHEK1 proteins and performed the kinase assay. Indeed, the N-terminus kinase domain had very high kinase activity, as reported by several studies (Caparelli and O'Connell, 2013; Emptage et al., 2017; Goto et al., 2015; Han et al., 2016; Tapia-Alveal et al., 2009; Walker et al., 2009). The R442Q mutant protein had about twice as high kinase activity as the WT protein (Figure 3A, Supplementary Figure 3). We next ask whether Chk1 R442Q may induce more DNA damage in zygotes. Interestingly, overexpression of Chk1 R442Q reduced γH2AX staining in the male pronucleus (Figure 3B and C, Supplementary Figure 4A). Since R442Q mutant protein has higher kinase activity than the WT protein, we used a CHEK1 inhibitor CCT244747 to treat zygotes overexpressing WT or R442Q. As expected, CCT244747 rescued the first mitosis arrest caused by R442Q, nearly all treated R442Q zygotes divided into 2-cell embryos (Supplementary Figure 4B). However, regular CCT244747 dosage also increased DNA damage, as shown by stronger γH2AX staining in treated WT or R442Q embryos (Supplementary Figure 4C and D). Since R442Q mutant has approximately double kinase activity than WT protein, we reasoned that reducing the kinase activity close to the normal level might be sufficient to overcome the first mitotic arrest. Indeed, 30 nM of CCT244747 permitted R442Q zygotes to divide into 2-cell embryos and did not increase the γH2AX signal (Figure 3D-G, Supplementary Figure 4D-F). Moreover, 60% of 30 nM CCT244747 treated R442Q zygotes developed to the blastocyst stage (Figure 3H and I). We examined the expression of pluripotency genes and found that Oct4 and Nanog proteins are highly expressed in the inner cell mass of CCT244747 treated R442Q blastocysts (Figure 3J). Treated WT and R442Q blastocysts also had a similar number of total cells and Oct4 and Nanog positive cells (Figure 3K), suggesting that their development was not compromised.

**Figure 3.**
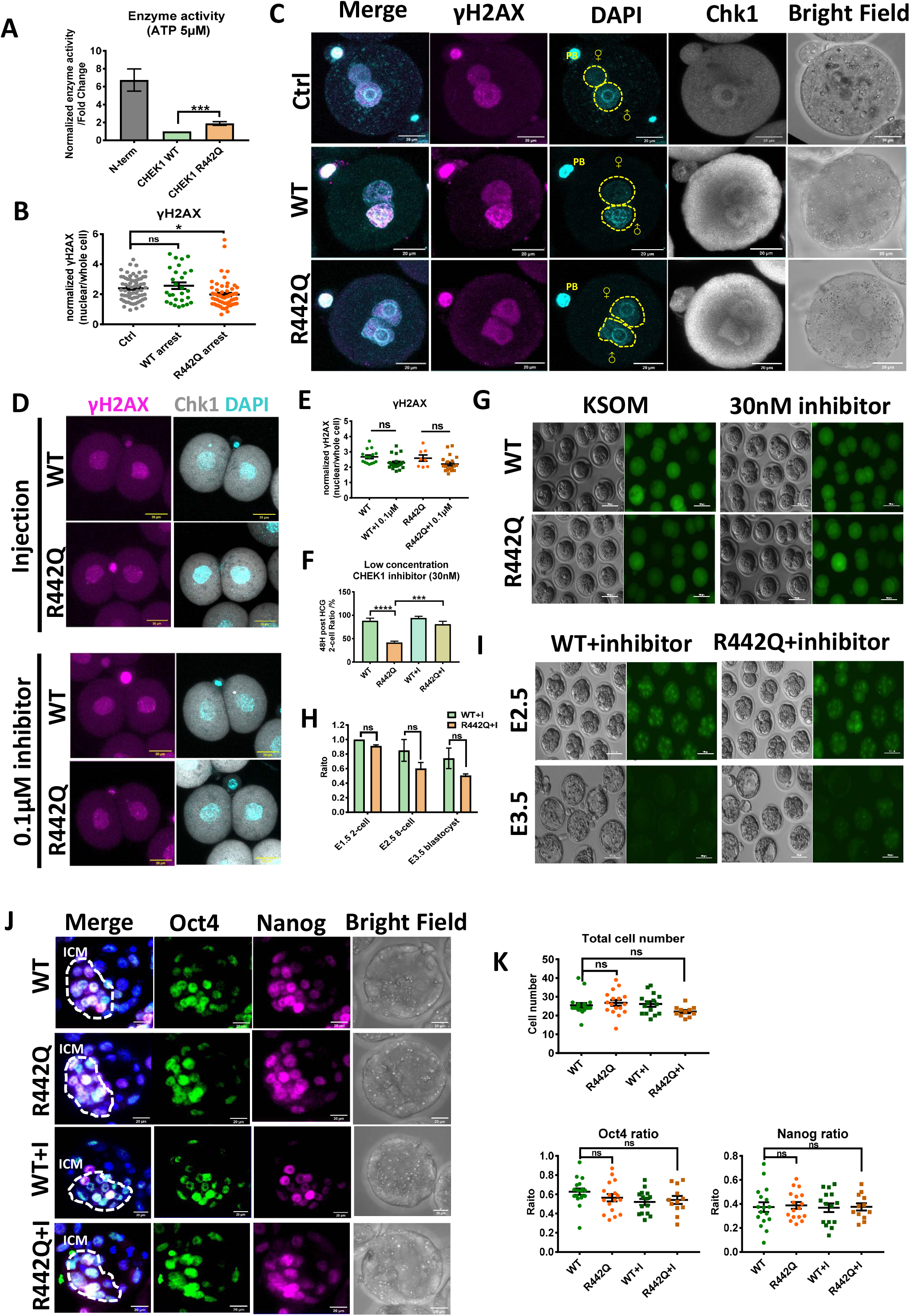
A graded Chk1 activity differentially modulates DNA damage response and cell cycle progression in the zygote. (A)CHEK1 R442Q protein showed higher kinase enzyme activity compared to WT. Luciferase based kinase assay normalized to WT group. 9 tests from 3 separate experiments. Two-sided Student's t-test. Columns are means ± s.e.m. (B)Dot plot quantification of γH2AX signal in arrested R442Q mouse zygotes compared to control non-injected (Ctrl) and WT injected zygotes. Ctrl, n=66, WT, n=28. R442Q, n=58. All pooled from 2 separate experiments. One-way analysis of variance (ANOVA), Bonferroni’s test for individual comparisons. Bars are means ± s.e.m. (C)Representative γH2AX immunostaining images of Ctrl, Chk1 WT, and R442Q zygotes. Yellow circles mark the paternal and maternal pronucleus. PB, polar body. Cyan, DAPI; pink, γH2AX; gray, Chk1. Scale bar, 20 μm. (D)γH2AX staining in CHEK1 inhibitor (0.1μM) rescued 2-cell embryos. Cyan, DAPI; pink, γH2AX; gray, Chk1. Scale bar, 20 μm. (E)Quantification of γH2AX signal in 0.1μM CHEK1 inhibitor-treated or non-treated WT and R442Q 2-cell embryos. Chk1 WT embryos with or without inhibitor treatment, n=15 and 21. Chk1 R442Q embryos with or without inhibitor treatment, n=8 and 20. One-way analysis of variance (ANOVA), Bonferroni’s test for individual comparisons. Bars are means ± s.e.m. (F)30 nM CHEK1 inhibitor treatment rescued the development of Chk1 R442Q zygotes. Chk1 WT embryos with or without 30nM inhibitor treatment, n=98 and 106. Chk1 R442Q embryos with or without 30nM inhibitor treatment, n=106 and 101. All pooled from 4 separated experiments. One-way analysis of variance (ANOVA), Bonferroni’s test for individual comparisons. Bars are means ± s.e.m. (G)Representative images of Chk1 WT and R442Q injected embryos with or without 30 nM inhibitor treatment at 48h post HCG. Scale bar, 100 μm. (H)30 nM CHEK1 inhibitor-treated embryos can develop to the blastocyst stage. Chk1 WT, n=48. Chk1 R442Q, n=46. All pooled from 2 separate experiments. Two-way analysis of variance (ANOVA), Bonferroni’s test for individual comparisons. Bars are means ± s.e.m. (I)Representative images of 8-cell and blastocysts developed from zygotes injected with mRNA of Chk1 WT and R442Q then treated with 30 nM CHEK1 inhibitor. The stage is as indicated. Scale bar, 100 μm. (J)Immunostaining of Oct4 and Nanog in blastocysts developed from WT or R442Q zygotes treated with 30 nM CHEK1 inhibitor. The dashed line highlights the ICM region. Blue, DAPI; green, Oct4; pink, Nanog. Scale bar, 20 μm. ICM, inner cell mass. (K)Statistic analysis of total cell number, the ratio of Oct4, and Nanog positive cells in blastocysts derived from Chk1 WT and R442Q injected zygotes with or without inhibitor treatment. WT embryos with or without 30nM inhibitor treatment, n=19 and 15. R442Q embryos with or without 30 nM inhibitor treatment, n=17 and 13. All pooled from 2 separate experiments. One-way analysis of variance (ANOVA), Bonferroni’s test for individual comparisons. Bars are means ± s.e.m.

We next performed transcriptome profiling of zygotes, 2-cell embryos, and blastocysts injected with mRNA encoding WT or R442Q with or without low dosage CHEK1 inhibitor treatment (Figure 4A). Principal component analysis (PCA) and heatmap clustering showed that 1-cell arrested R442Q zygotes clustered together with PN5 WT zygotes (Figure 4B and C), suggesting that the minor zygotic genome activation was not affected. At the late 2-cell stage, the transcriptome of R442Q embryos, which managed to divide, clustered together with WT 2-cell embryos (Figure 4B and C), indicating that the major zygotic genome activation occurred in these embryos. The CHEK1 inhibitor rescued WT and R442Q blastocysts clustered together, consistent with our previous observations (Figure 4B and C, Figure 3J and K). We analyzed differentially expressed transcripts in WT and R442Q zygotes and 2-cell embryos. Compare to WT embryos, R442Q zygotes downregulated RNA processing and cell division related genes. R442Q 2-cell embryos decreased the expression of some RNA processing and chromatin modification genes but increased genes associated with nucleotide metabolism (Supplementary Figure 5). Finally, we transferred CHEK1 inhibitor rescued R442Q 2-cell embryos into female mice and found that a similar number of pups were born compared with the WT embryo transferred group (Figure 4D and E, Supplementary Figure 3G). These results suggest that the low concentration of CHEK1 inhibitor treatment may relieve zygotes from the first mitotic division arrest caused by a slight tightening up of DNA damage checkpoint, and the rescued embryos would develop normally.

**Figure 4.**
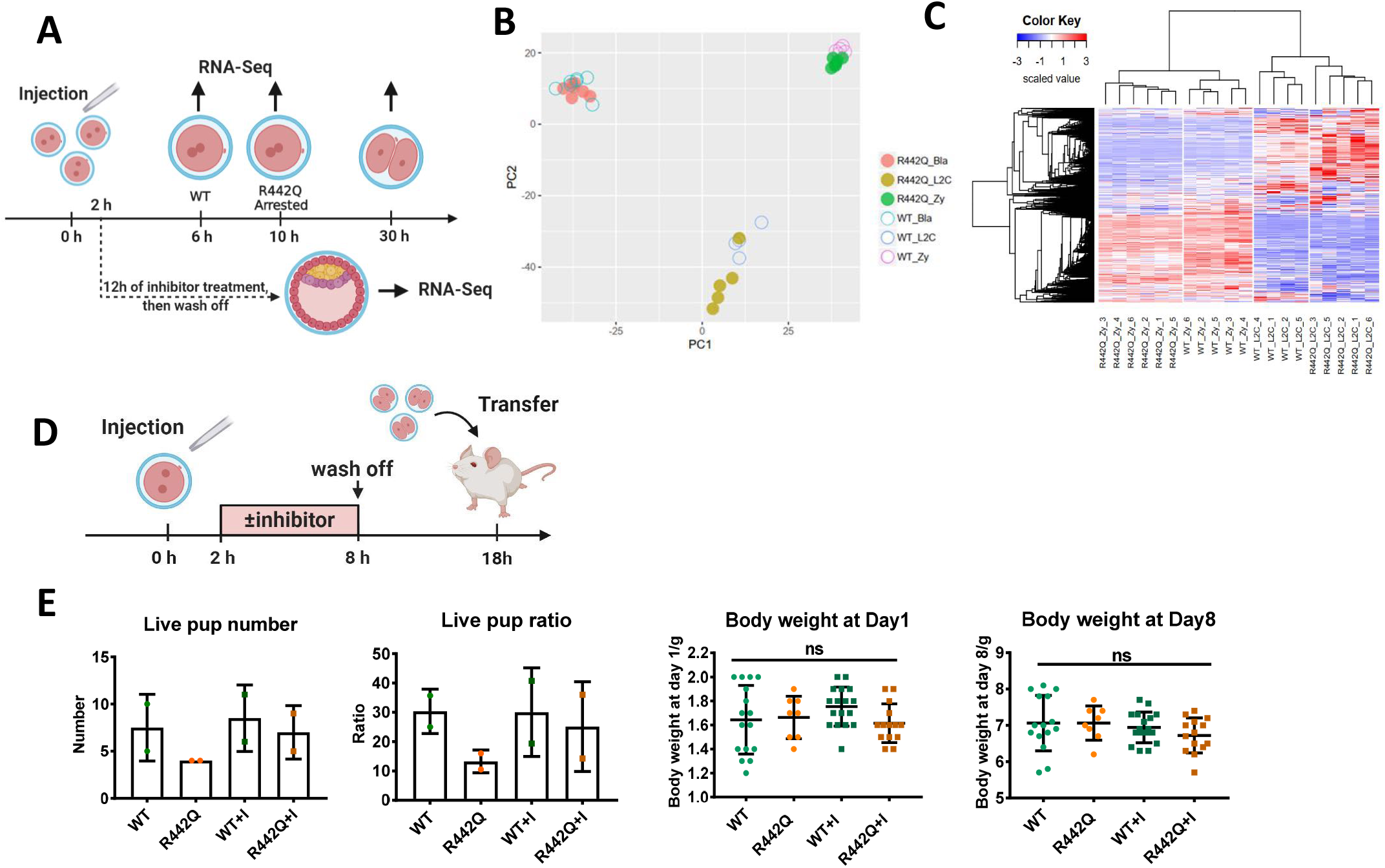
A low dosage of CHEK1 inhibitor treatment does not affect zygotic genome activation and permitted normal embryo development. (A)Schematic of Chk1 WT and R442Q embryos used for RNA-Seq analysis (Created with BioRender.com). Chk1 WT or R442Q mRNA were injected into mouse zygotes. One-cell embryos were harvested at 6h (WT) or 10h (R442Q) post-injection. Two-cell embryos were collected 30h post-injection. For blastocysts, Chk1 WT and R442Q injected zygotes were first treated with 30nM inhibitor overnight. After washing off the inhibitor, 2-cell embryos were cultured to the blastocyst stage and used for RNA-seq. (B)PCA analysis of RNA-Seq data from single WT and R442Q embryos. (C)Heatmap clustering of WT and R442Q zygotes and 2-cell stage embryos. (D)Schematic of Chk1 WT and R442Q injected zygotes with or without 30 nM of CHEK1 inhibitor treatment, followed by transferring to pseudopregnant female mice (Created with BioRender.com). (E)Chk1 WT and R442Q embryos treated with or without 30 nM of inhibitor developed to live pups with normal body weight after transplantation. n=15, 8, 17, and 14 live pups of Chk1 WT, R442Q, WT with inhibitor, and R442Q with inhibitor group. All pooled from 2 separate experiments.

The discovery of the *CHEK1* R442Q mutation and the first zygotic mitosis arrest phenotype in human reproduction has important implications. All the heterozygous female carriers are infertile, presumably due to failed mitosis in zygotes. As the zygotes contained mostly maternal proteins inherited from the egg, both CHEK1 WT and R442Q proteins should be present in the carrier's eggs. Based on the clinical observation during the IVF treatment, CHEK1 R442Q did not affect oocyte meiosis, suggesting that a small elevation in CHEK1 kinase activity does not block the first meiotic division in women. The II-8 and III-8 male carriers are healthy and do not have fertility problems, indicating that *CHEK1* R442Q does not affect sperm meiosis in men. CHEK1 is a ubiquitously expressed protein, and all the heterozygous individuals are healthy without cancer or autoimmune diseases based on a consent health history survey. Thus, CHEK1 R442Q should not affect most mitotic division of embryonic or adult cells. All the heterozygous females inherited the *CHEK1* R442Q mutation from their father as their mother has two copies of the normal *CHEK1* gene. Since the paternal genome starts to express after the major zygotic genome activation during the 4 to 8-cell stage in humans (Kermi et al., 2019; Khokhlova et al., 2020; Xue et al., 2013; Yan et al., 2013), the CHEK1 R442Q protein derived from the mutated paternal copy of *CHEK1* will likely be present from the 8-cell stage onwards. Therefore, the small elevation of CHEK1 kinase activity does not appear to arrest cell division after the 8-cell stage in human embryos. Thus, the CHEK1 R442Q protein seemed to affect the first mitotic division most seriously, indicating that the first mitosis is particularly sensitive to DNA damage checkpoint activation in humans. Moreover, the DNA damage checkpoint threshold may be cell type, developmental stage, and species-dependent, based on the human phenotype and our results obtained in the mouse system. When we treat Chk1 R442Q overexpressing mouse zygotes with a low dosage of CHEK1 inhibitor, they can escape the first mitotic arrest and develop normally. CHEK1 inhibitors are often used as cancer drugs. They inhibit the DNA damage checkpoint, making cancer cells accumulate DNA damage and die (Dent et al., 2011; Goto et al., 2012; McNeely et al., 2014; Smith et al., 2010; Zhang and Hunter, 2014). Surprisingly, although a high dosage of CHEK1 inhibitor also induced DNA damage in zygotes, a low dosage of CHEK1 inhibitor rescued R442Q overexpression induced first mitosis arrest without eliciting any DNA damage judging by the γH2AX staining, and the embryos developed normally. Chk1 R442Q overexpression also did not induce γH2AX signal. The above results suggested that small up or down-regulation of CHK1 kinase activity may not affect genome stability. Our findings also imply the possibility to use low dosage CHEK1 inhibitor treatment to release cells from low-level genome stress induced cell cycle block during zygote mitosis or other sensitive periods.

## METHODS

### Patient consent and ethical approvement

The proband III-7 and her family were recruited for this study from the First Affiliated Hospital of Anhui Medical University. The human sample collection and analysis was approved by the Ethics Committee for Clinical Medical Research of the First Affiliated Hospital of Anhui Medical University and carried out with informed consent.

### Whole-exome sequencing (WES) analysis

Libraries were generated using the Agilent SureSelect Human All Exon V6 kit (Agilent Technologies, USA) following the manufacturer’s recommendations. WES was carried out on an Illumina NovaSeq6000 sequencer with pair end 150 bp (PE150) for each reaction. The original sequencing reads were aligned to the reference genome GRCh37/hg19, variants including single nucleotide polymorphisms (SNPs) and short insertion and deletion (INDELs) were called using the GATK software program. The identified variants were further annotated using ANNOVAR software. The criteria used for filtering the desired variants were as follows (i) missense, nonsense, frameshift, or splice site variants; (ii) variants with minor allele frequency < 1%. The minor allele frequency data were obtained by referring to the following databases: Genome Aggregation Database (gnomAD, http://gnomad.broadinstitute.org/), 1000 Genomes (1000G, http://browser.1000genomes.org/index.html), and the NHLBI Exome Sequencing Project (ESP6500).

### Quantitative PCR

Total RNA of patient and control blood samples were extracted with TRIZOL (Invitrogen). 1μg RNA of each sample was used for reverse transcription with Superscript II (Invitrogen). Q-PCR reactions were performed using GoTaq qPCR Master Mix (Promega) in a CFX96 Real-Time System (Bio-Rad). The relative expression level of each gene was normalized against the Ct (Critical Threshold) value of the house-keeping gene GAPDH using the Bio-Rad CFX Manager program. For detecting the mutation transcript in the patient’s somatic cells, qPCR products were purified by agarose gel electrophoresis, ligated to pEASY-T5 Zero plasmid, and sequenced using the M13 forward primer.

### Mouse embryo culture and microinjection

All animal experiments were conducted following the Guide for the Care and Use of Animals for Research Purposes. The protocol for mouse embryo isolation was approved by the Institutional Animal Care and Use Committee and Internal Review Board of Tsinghua University. Embryos were collected from wild type C57BL/6 females (Charles River). Zygotes for RNA injection were collected from mated female mice 26-28 hrs post-HCG, then cultured in KSOM medium in 37.5℃ 5% CO2 incubator. GV oocytes were collected from the ovary of C57BL/6 female in M2 medium supplemented with 10μM Cilostamide. Microinjection of mRNAs at a concentration of 200 ng/μl into the mouse GV oocytes or PN3 zygotes was performed on a Leica microscope micromanipulator. The microinjected embryos were treated with or without CHEK1 inhibitor CCT244747 at different concentrations as indicated.

### Plasmid cloning and RNA synthesis

Mouse *Chk1* cDNA was cloned into the RN3P vector for in vitro transcription of mRNA. mRNAs were generated using the T3 mMESSAGE mMACHINE Kit according to the manufactures instructions (Ambion).

### CHEK1 protein production and kinase assay

For human CHEK1 protein purification and kinase assay, N-term (1-289aa), WT full length, and mutant full-length human *CHEK1* cDNAs were cloned from 293FT cells, fused with 6×His tag at the N-terminus, and cloned into Bac-to-Bac Baculovirus Expression System following the instructions of the manufactures.

For kinase assay, Sf9 insect cells cultured in suspension in ESF 921 Insect Cell Culture Medium were used for protein overexpression and purification. Baculovirus of His-tagged N-term (1-289aa), WT full length, and mutant full-length human *CHEK1* were introduced into the sf9 insect cells by Cellfectin II Reagent following the instructions of the manufactures. Recombinant Proteins were purified with affinity Ni-NTA agarose. Purified proteins were verified by Coomassie bright blue stain and Western blot with the antibody of CHEK1 and His-tag. CHEK1 kinase assay was conducted using the kinase assay kit with the substrate of Cdc25C-derived Chktide.

### Immunostaining and confocal microscopy

For immunostaining, mouse oocytes or embryos were fixed, then permeabilized with Triton X-100 for 20 min. Embryos were then blocked with 5% FCS at room temperature for 2 h and incubated with primary antibody overnight at 4 ℃,followed by washing in PBST and incubating with secondary antibody for 1 h at room temperature. Immunostaining images were acquired with a Nikon A1 confocal microscope using an oil-immersion 40× objective. Raw data were processed using open-source image analysis software Fiji ImageJ.

### Transcriptome Profiling

For gene expression study in mouse preimplantation embryos, we used the SMART-seq2 protocol to amplify single embryo RNA (Picelli et al., 2014). Embryos were first lysed in hypotonic lysis buffer, followed by reverse transcription and pre-amplification. After AMPure XP beads purification, the cDNA libraries were tagmented by Tn5 and were subject to Illumina Nextera library preparation. All libraries were sequenced on Illumina HiSeq X-10 according to the manufacturer’s instruction.

### RNA-seq data processing

Adapter sequences were trimmed using TrimGalore (version 0.4.4). Clean reads were mapped to the mouse genome (mm10) using Bowtie2 (version 2.3.5) software with the Refseq annotation. Gene expression values were represented with FPKM (Fragments Per Kilobase per Million) calculated by RSEM (version 1.2.28). Differential expression genes (DEGs) were analyzed using the DESeq2 package (version 1.20.0). Genes that satisfied the threshold “fold change ≥ 2, p < 0.01” were considered significant DEGs. Principle component analyses were performed using R's “prcomp” function and drawn by ggplot2 package (version 3.1.0). The heatmaps were produced by the heatmap2 function of the gplots package (version3.0.1.1) with the hierarchical clustering method. The enrichment of GO was analyzed using clusterProfiler package (version 3.8.1).

### Statistical analysis

Data are presented as mean ± standard error of the mean (SEM). Statistical significance was determined by Student’s t-test (two-tail) for two groups; one-way or two-way Analysis of Variance (ANOVA) for multiple groups using Graphpad software. P < 0.05 was considered significant.

## AUTHOR CONTRIBUTIONS

BC, JG, TW, QL, ZZ, LL, YC, and JN conceived the study and designed the experiments. JG, TW, QL, JM performed mouse embryo and oocytes collection and microinjection, phenotype analysis, and transcriptome profiling; BC and FD performed clinical sample collection and data analysis, HL instructed protein kinase activity study, JG, BC, TW, ZZ, LL, YC and JN wrote the manuscript. All authors read and approved the final version of the manuscript.

## COMPETING INTEREST STATEMENT

The authors declare no competing interests.

## Data availability

Raw and processed RNA-seq data are available at the Gene Expression Omnibus GEO.

---

**Supplemental figure 1.**
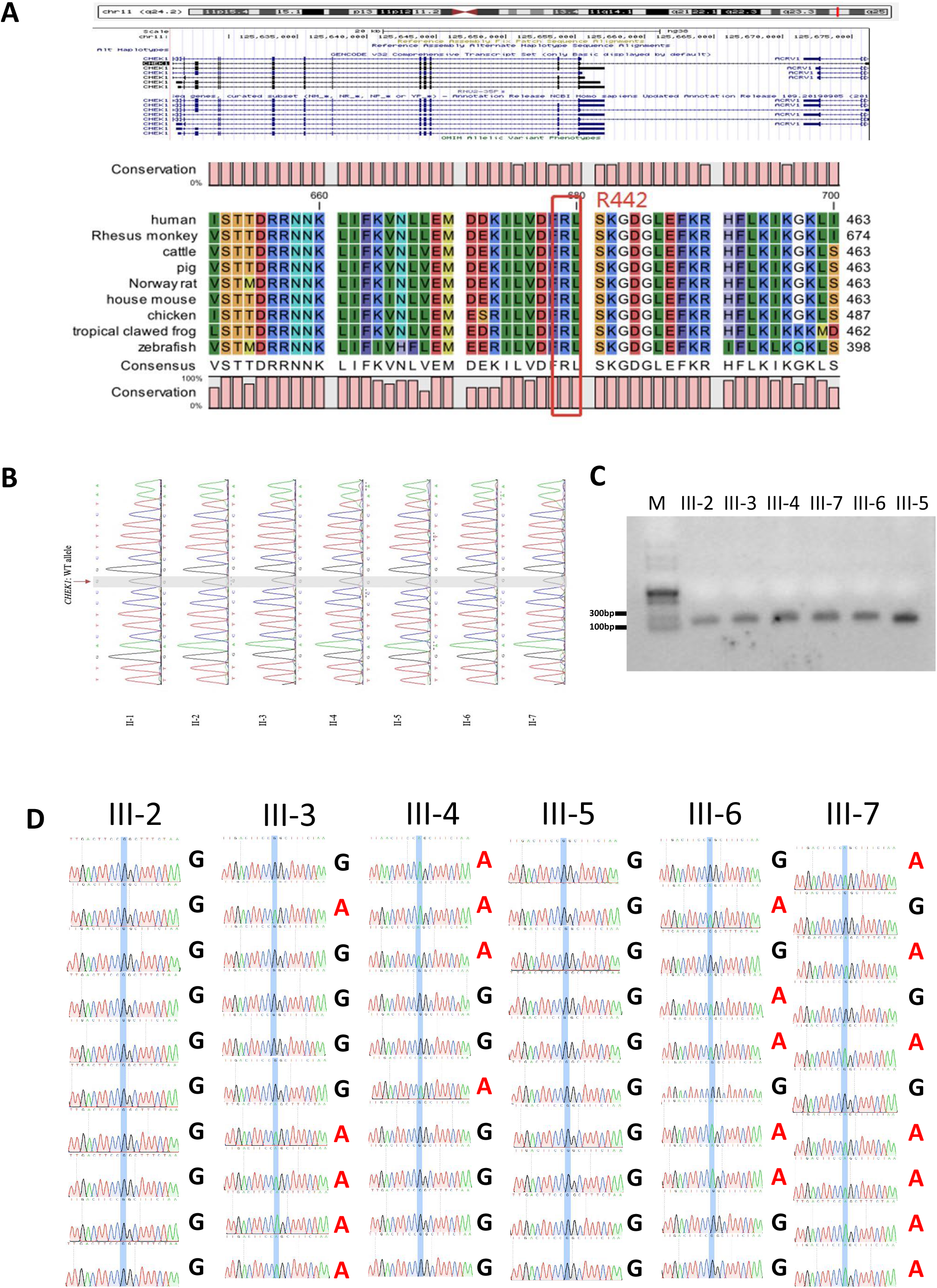
*CHEK1* gene locus and *CHEK1* R442Q expression in somatic cells of carriers. (A)Human *CHEK1* gene locus and amino acid sequence of CHEK1 protein C-terminus. The R442 in human CHEK1 protein is conserved among all species. (B)No *CHEK1* R442Q mutation was detected by Sanger sequencing in patients’ father’s siblings. Single peak of G read indicates the normal *CHEK1* allele. (C)*CHEK1* is expressed in somatic cells of carriers verified by RT-PCR and agarose gel electrophoresis. (D)Detection of the ratio of *CHEK1* R442Q mutant RNA transcript by Sanger sequencing. 10 colonies were picked and sequenced for each sample.

**Supplemental figure 2.**
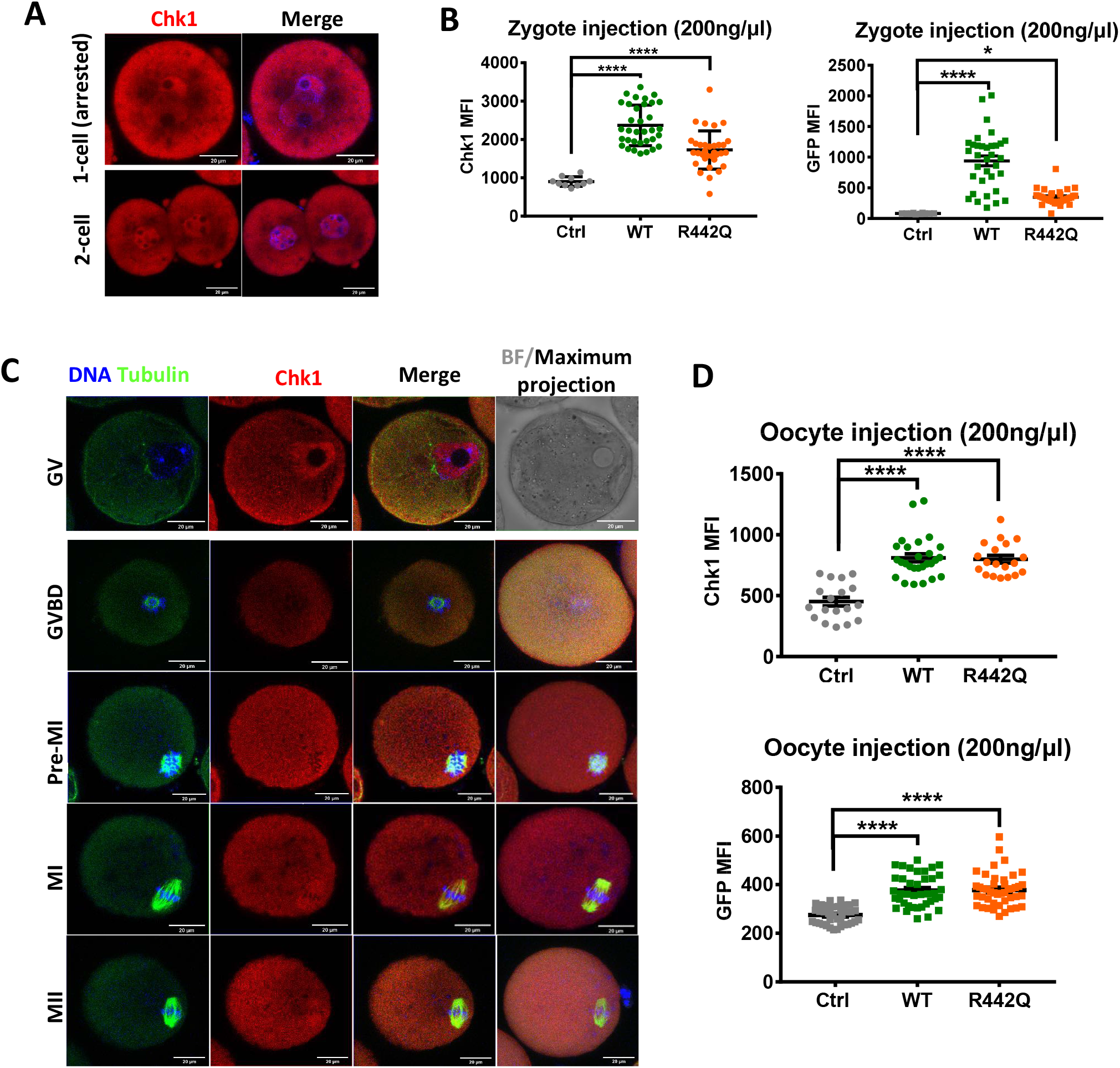
Chk1 localization in mouse oocytes and preimplantation embryos. (A)Chk1 protein localizes to both nuclear and cytoplasm in mouse early embryos. Red, Chk1; blue, DAPI. Scale bar, 20 μm. (B)Microinjection of mRNA encoding Chk1 WT and R442Q GFP fusion protein leads to an elevation of Chk1 and GFP protein expression in zygotes and 2-cell embryos. Ctrl, n=10. WT, n=34. R442Q, n=33. All pooled from 2 separate experiments. One-way analysis of variance (ANOVA), Bonferroni’s test for individual comparisons. Bars are means ± s.e.m. (C)Chk1 protein locates in the germinal vesicle and on the spindle of mouse oocytes. Red, Chk1; blue, DAPI; green, Tubulin. Scale bar, 20 μm. (D)Microinjection of mRNA encoding Chk1 WT and R442Q GFP fusion protein into GV oocytes leads to an elevation of Chk1 and GFP protein expression. Ctrl, n=38. WT, n=39. R442Q, n=44. All pooled from 2 independent experiments. One-way analysis of variance (ANOVA), Bonferroni’s test for individual comparisons. Bars are means ± s.e.m.

**Supplemental figure 3.**
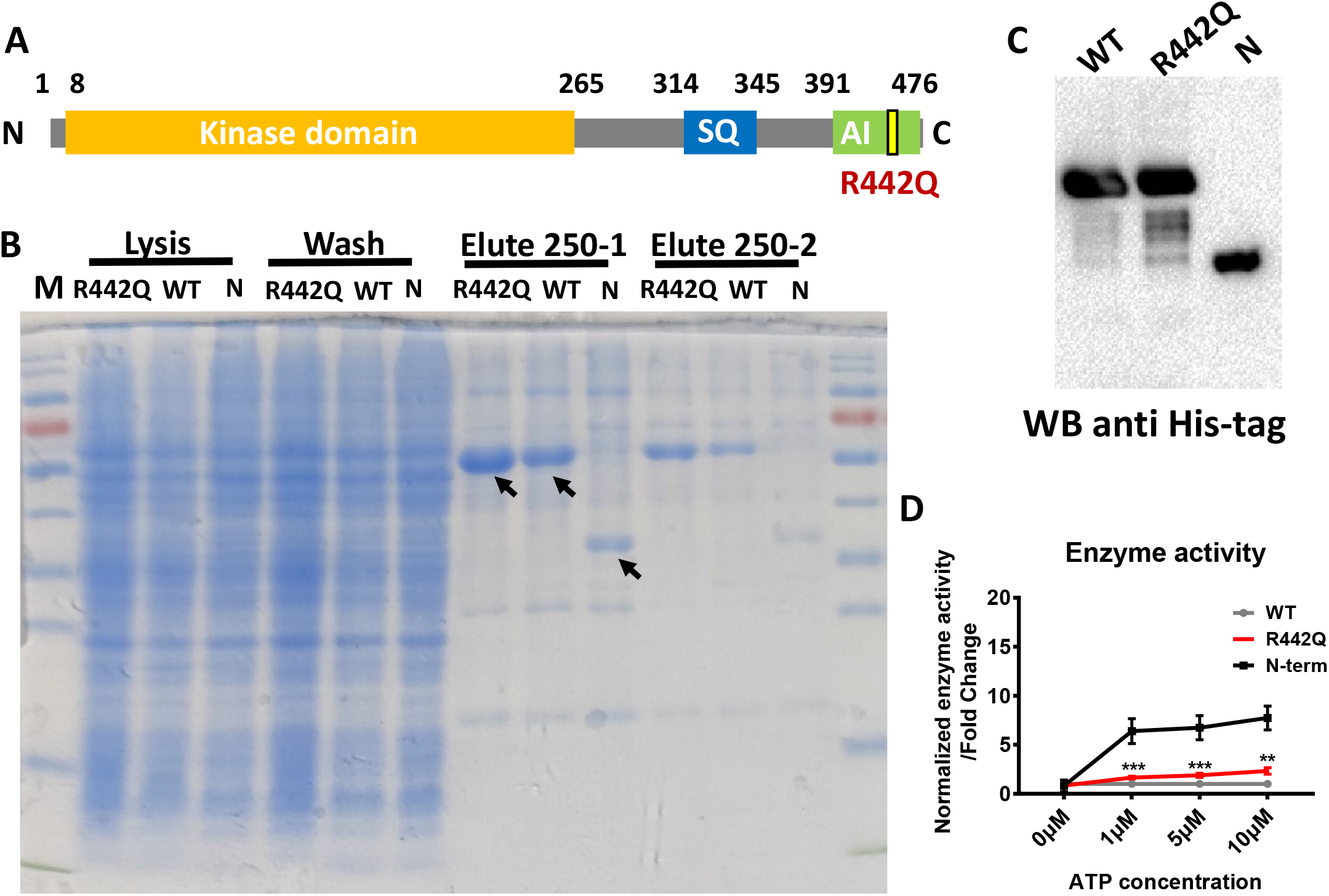
CHEK1 R442Q showed an elevated kinase activity compared to WT. (A)Schematic of human CHEK1 protein domains. (B)Coomassie blue staining showing purification of CHEK1 N-term, WT, and R442Q proteins. M, marker. Arrow points to the purified His-tagged proteins. (C)Western blot showing His-tag purified CHEK1 N-term, WT, and R442Q proteins. (D)Kinase assay showed CHEK1 R442Q mutant protein had increased kinase activity compared to WT protein. n=9 from 3 independent experiments. Two-sided Student's t-test. Columns are means ± s.e.m.

**Supplemental figure 4.**
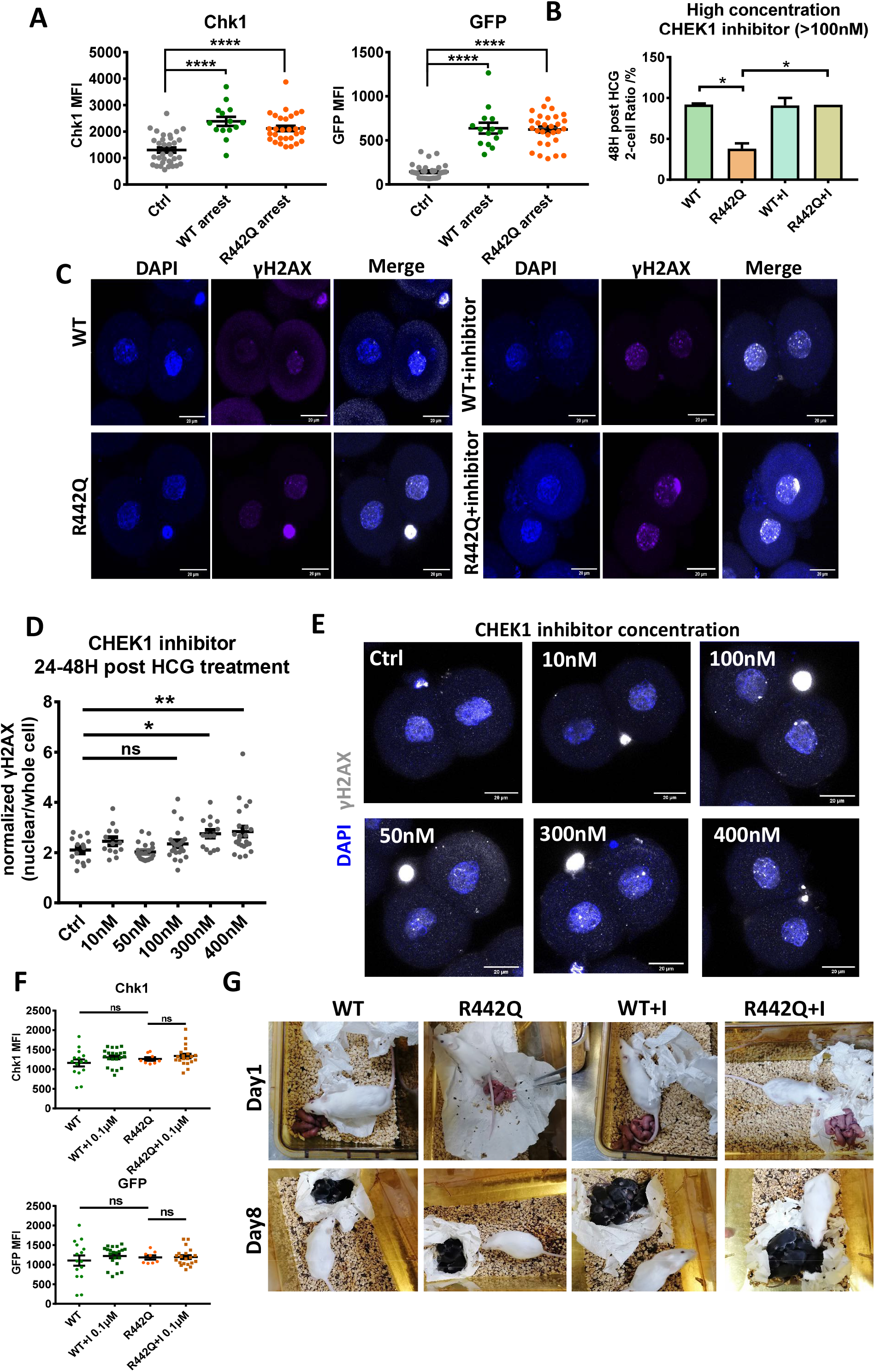
A high concentration of CHEK1 inhibitor treatment caused DNA damage. (A)Quantification of Chk1 and GFP protein levels in zygotes injected with mRNA encoding Chk1 WT and R442Q GFP fusion proteins. Related to Figure 3C. n=66 Ctrl zygotes, n=28 WT oocytes, and n=58 R442Q oocytes, pooled from 2 separate experiments. One-way analysis of variance (ANOVA), Bonferroni’s test for individual comparisons. Bars are means ± s.e.m. (B)High concentration of CHEK1 inhibitor (>100 nM) rescue the Chk1 R442Q injected embryos to the 2-cell stage at 48h post HCG. Chk1 WT injected embryos with or without inhibitor treatment, n=41 and 29; Chk1 R442Q injected embryos with or without inhibitor treatment, n=44 and 24. All pooled from 2 separate experiments. One-way analysis of variance (ANOVA), Bonferroni’s test for individual comparisons. Bars are means ± s.e.m. (C)High concentration of CHEK1 inhibitor (500 nM) leads to increased DNA damage response in WT and R442Q 2-cell embryos. Blue, DAPI; purple, γH2AX. Scale bar, 20 μm. (D)CHEK1 inhibitor concentration lower than 100 nM did not increase DNA damage response in mouse early embryos. n=15,13,21,22,15 and 23 of Ctrl, 10 nM, 50 nM, 100 nM, 300 nM and 400 nM inhibitor-treated groups. One-way analysis of variance (ANOVA), Bonferroni‘s test for individual comparisons. Bars are means ± s.e.m. (E)Representative images of mouse embryos treated with different concentrations of CHEK1 inhibitor 48h post HCG from (D). Blue, DAPI; gray, γH2AX. Scale bar, 20 μm. (F)Comparing Chk1 and GFP protein levels in 2-cell embryos injected with mRNA encoding Chk1 WT and R442Q GFP fusion protein at PN3 zygote stage and treated with or without 0.1 µM of CHEK1 inhibitor, related to Figure 3D. WT embryos with or without inhibitor treatment, n=15 and 21. R442Q embryos with or without inhibitor treatment, n=8 and 20. All pooled from 2 separate experiments. One-way analysis of variance (ANOVA), Bonferroni’s test for individual comparisons. Bars are means ± s.e.m. (G)Chk1 WT and R442Q mouse embryos treated with 30 nM of inhibitor developed to live pups with normal body weight after transplantation. Related to Figure 4E.

**Supplementary figure 5.**
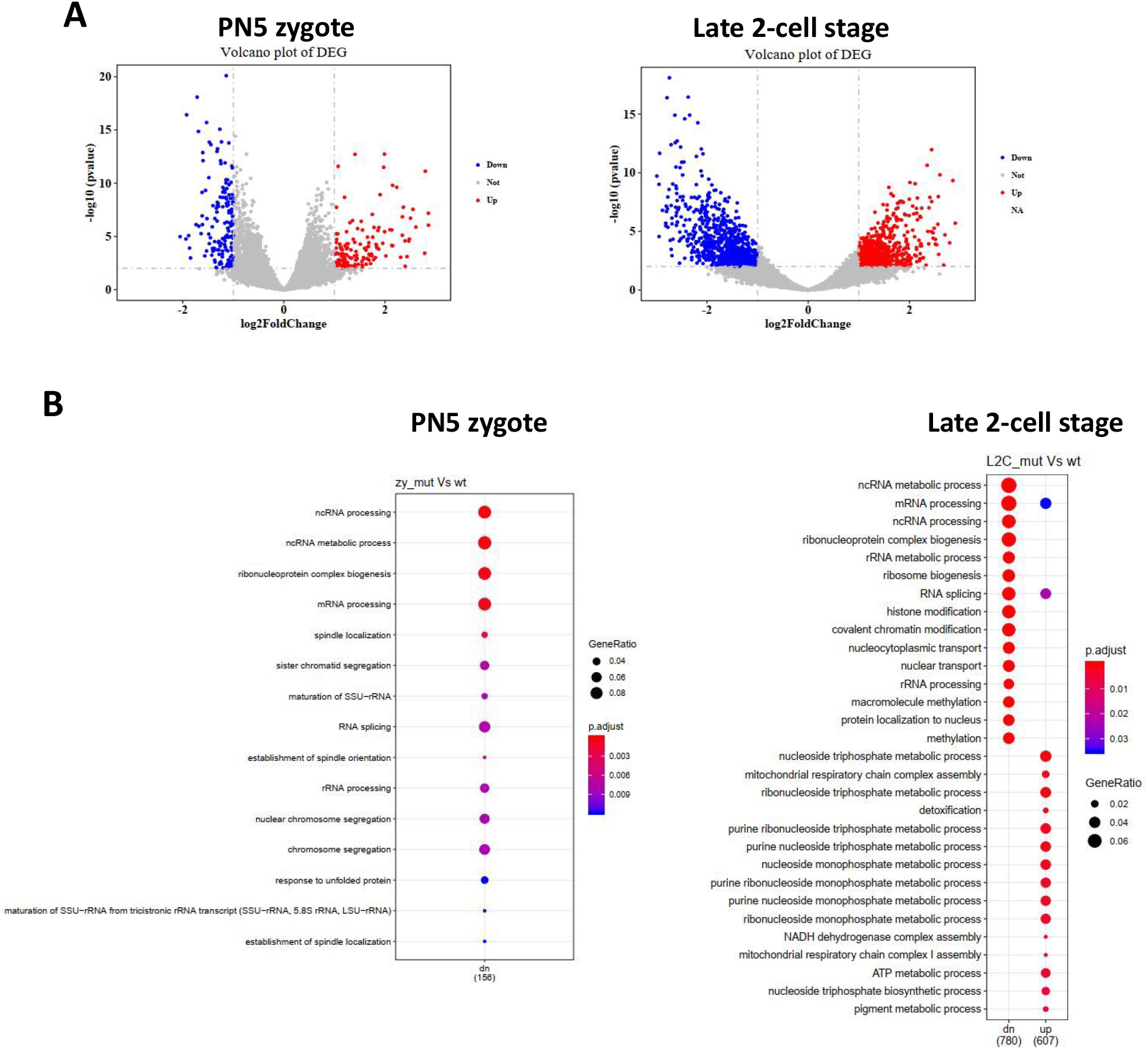
Transcriptome analysis of mouse embryos injected with mRNA of Chk1 WT and R442Q. (A)Volcano plot depicting significantly differentially expressed genes in zygotes and 2-cell embryos injected with mRNA of Chk1 WT (n=5 zygotes, n=4 2-cell embryos) and R442Q (n=6 zygotes, n=5 2-cell embryos). (B)Bubble plots showing the major GO terms of differentially expressed genes enriched in Chk1 WT and R442Q zygotes and late 2-cell embryos.

## Notes

### Competing Interest Statement

The authors have declared no competing interest.

### Summary of Updates

Anyone who wishes to share, reuse, remix, or adapt this material must obtain permission from the corresponding author.

## References

Caparelli, M.L., and O'Connell, M.J. (2013). Regulatory motifs in Chk1. Cell Cycle 12, 916–922.

Chen, L., Chao, S.B., Wang, Z.B., Qi, S.T., Zhu, X.L., Yang, S.W., Yang, C.R., Zhang, Q.H., Ouyang, Y.C., Hou, Y., et al. (2012). Checkpoint kinase 1 is essential for meiotic cell cycle regulation in mouse oocytes. Cell Cycle 11, 1948–1955.

Clift, D., and Schuh, M. (2013). Restarting life: fertilization and the transition from meiosis to mitosis. Nature Reviews Molecular Cell Biology 14, 549–562.

Dent, P., Tang, Y., Yacoub, A., Dai, Y., Fisher, P.B., and Grant, S. (2011). CHK1 Inhibitors in Combination Chemotherapy Thinking Beyond the Cell Cycle. Mol Interv 11, 133–140.

Emptage, R.P., Schoenberger, M.J., Ferguson, K.M., and Marmorstein, R. (2017). Intramolecular autoinhibition of checkpoint kinase 1 is mediated by conserved basic motifs of the C-terminal kinase-associated 1 domain. Journal of Biological Chemistry 292, 19024–19033.

Goto, H., Izawa, I., Li, P., and Inagaki, M. (2012). Novel regulation of checkpoint kinase 1: Is checkpoint kinase 1 a good candidate for anti-cancer therapy? Cancer Sci 103, 1195–1200.

Goto, H., Kasahara, K., and Inagaki, M. (2015). Novel Insights into Chk1 Regulation by Phosphorylation. Cell Struct Funct 40, 43–50.

Han, X.Z., Tang, J.S., Wang, J.N., Ren, F., Zheng, J.H., Gragg, M., Kiser, P., Park, P.S.H., Palczewski, K., Yao, X.S., et al. (2016). Conformational Change of Human Checkpoint Kinase 1 (Chk1) Induced by DNA Damage. Journal of Biological Chemistry 291, 12951–12959.

Ju, J.Q., Li, X.H., Pan, M.H., Xu, Y., Sun, M.H., Xu, Y., and Sun, S.C. (2020). CHK1 monitors spindle assembly checkpoint and DNA damage repair during the first cleavage of mouse early embryos. Cell Proliferat 53.

Kermi, C., Aze, A., and Maiorano, D. (2019). Preserving Genome Integrity during the Early Embryonic DNA Replication Cycles. Genes-Basel 10.

Khokhlova, E.V., Fesenko, Z.S., Sopova, J.V., and Leonova, E.I. (2020). Features of DNA Repair in the Early Stages of Mammalian Embryonic Development. Genes-Basel 11.

Liu, Q.H., Guntuku, S., Cui, X.S., Matsuoka, S., Cortez, D., Tamai, K., Luo, G.B., Carattini-Rivera, S., DeMayo, F., Bradley, A., et al. (2000). Chk1 is an essential kinase that is regulated by Atr and required for the G(2)/M DNA damage checkpoint. Gene Dev 14, 1448–1459.

McNeely, S., Beckmann, R., and Lin, A.K.B. (2014). CHEK again: Revisiting the development of CHK1 inhibitors for cancer therapy. Pharmacol Therapeut 142, 1–10.

Patil, M., Pabla, N., and Dong, Z. (2013). Checkpoint kinase 1 in DNA damage response and cell cycle regulation. Cellular and Molecular Life Sciences 70, 4009–4021.

Picelli, S., Faridani, O.R., Bjorklund, A.K., Winberg, G., Sagasser, S., and Sandberg, R. (2014). Full-length RNA-seq from single cells using Smart-seq2. Nat Protoc 9, 171–181.

Smith, J., Tho, L.M., Xu, N.H., and Gillespie, D.A. (2010). The ATM-Chk2 and ATR-Chk1 Pathways in DNA Damage Signaling and Cancer. Adv Cancer Res 108, 73–112.

Tapia-Alveal, C., Calonge, T.M., and O'Connell, M.J. (2009). Regulation of Chk1. Cell Div 4.

Walker, M., Black, E.J., Oehler, V., Gillespie, D.A., and Scott, M.T. (2009). Chk1 C-terminal regulatory phosphorylation mediates checkpoint activation by de-repression of Chk1 catalytic activity. Oncogene 28, 2314–2323.

Xue, L., Cai, J.Y., Ma, J., Huang, Z., Guo, M.X., Fu, L.Z., Shi, Y.B., and Li, W.X. (2013). Global expression profiling reveals genetic programs underlying the developmental divergence between mouse and human embryogenesis. Bmc Genomics 14.

Yan, L.Y., Yang, M.Y., Guo, H.S., Yang, L., Wu, J., Li, R., Liu, P., Lian, Y., Zheng, X.Y., Yan, J., et al. (2013). Single-cell RNA-Seq profiling of human preimplantation embryos and embryonic stem cells. Nat Struct Mol Biol 20, 1131–+.

Zhang, Y.W., and Hunter, T. (2014). Roles of Chk1 in cell biology and cancer therapy. Int J Cancer 134, 1013–1023.

